# Managing invasive hybrids through habitat restoration in an endangered salamander system

**DOI:** 10.1101/2022.11.09.515819

**Authors:** Robert D. Cooper, H. Bradley Shaffer

**Affiliations:** Department of Ecology and Evolutionary Biology, University of California, Los Angeles, CA 90095, USA; La Kretz Center for California Conservation Science, Institute of the Environment and Sustainability, University of California, Los Angeles, CA 90095, USA

**Keywords:** Hybridization, Invasive species, Conservation, *Ambystoma californiense*, Hydroperiod

## Abstract

Invasive species present one of the greatest threats to the conservation of biodiversity. When invasives hybridize with endangered native taxa, they introduce novel challenges ranging from the identification of hybrids in the field, to hybrid vigor and the erosion of species identity as genotypes are lost. Across a large swath of central California, a hybrid swarm consisting of admixed endangered California tiger salamanders (“CTS”, *Ambystoma californiense*) and introduced barred tiger salamander (*Ambystoma mavortium*) has replaced native populations, threatening CTS with genomic extinction. Here we employ a large-scale, genomically-informed field ecological experiment to test whether habitat restoration can reinstate natural selection favoring native salamander genotypes. We constructed 14 large, semi-natural ponds and manipulated their hydroperiods to evaluate larval survival and mass at metamorphosis. Consistent with earlier work, we found overwhelming evidence of hybrid superiority which persisted across all hydroperiod treatments. Short duration ponds substantially reduced the mass and survival probability of both native and hybrid larvae, likely exerting strong selective pressure in the wild. We identified 86 candidate genes, representing 1.8% of 4,723 screened loci, that significantly responded to this hydroperiod-driven selection. In contrast to previous mesocosm-based studies, native CTS never exhibited greater fitness than hybrids, suggesting that hydroperiod management alone will not shift selection to favor native genotypes. However, shortening pond hydroperiod may represent a cost-effective strategy to limit the overall productivity of ponds with non-native genotypes, complimenting additional strategies such as targeted hybrid removal. At a broader level, our experimental approach leverages extensive ecological knowledge, modern genomic tools, and a naturalistic, *in situ* replicated design to critically evaluate and expand the potential toolkit that managers can use to address this, and other recalcitrant biological invasions. We believe that this strategy may be an important tool for managing the growing number of complex invasion scenarios threatening global biodiversity.

## Introduction

The introduction and establishment of invasive taxa represents one of the most challenging conservation concerns of the 21^st^ century (1–3). Though most introduced populations do not persist, those that do can cause tremendous harm to ecosystem stability (4), public health (5) and local economies, costing the US alone an estimated $100 billion dollars annually (6). Attempts at wholesale eradications to remove invasive species (7, 8) can be labor-intensive and involve billions of dollars and monumental human effort to manually eradicate unwanted species (9). Though occasionally effective (10), eradication is often not the optimal solution (11, 12), especially for species that are difficult to find or closely resemble native taxa. Eradication is both a challenging and potentially harmful endeavor (13), and is particularly problematic if their native counterparts are rare or endangered.

An appealing alternative strategy for managing biological invasions leverages natural selection, enabled through habitat restoration, to favor native taxa and reduce or eliminate non-native individuals or genotypes. Human-mediated habitat degradation is one of the most common predictors of successful biological invasions (14, 15). Such landscape modification can shift selective pressures, creating novel, invadable niches that select for non-native genotypes (e.g. 16, 17). Recent work demonstrating that healthy, intact ecosystems are resistant to biological invasions suggests that many can be slowed, or possibly reversed, by restoring habitat features to reinstate the natural selection pressures that shaped the evolution of native communities (18, 19). Habitat restoration is a particularly attractive tool for invasive species management when the life history or behavioral ecology of a species renders direct eradication difficult or impossible (20, 21). While intuitively appealing, few empirical studies have directly tested the efficacy of habitat restoration as an invasive species control measure, likely due to logistical constraints involved in rigorously testing this idea. The scale required to statistically examine the effects of habitat modification on target species makes this research program prohibitive for many species, despite the growing demand for effective, ecologically driven invasive species control programs.

In this study, we evaluate what is perhaps the key component of vernal pool habitat restoration, pond hydroperiod, as a potential strategy to control, and possibly eradicate, non-native hybrids in an endangered amphibian ecosystem. The California tiger salamander (*Ambystoma californiense*; hereafter “CTS”) is an endemic California species restricted to vernal pool grassland habitats in central California (22). As aquatic larvae, CTS are important apex predators that help shape vernal pool communities and trophic webs (23–25). This endangered species is protected at both the federal and state levels based on population declines throughout its range (26, 27). One of the most challenging issues impeding CTS recovery is hybridization with introduced populations of the non-native barred tiger salamander (*Ambytoma mavortium*, hereafter “BTS”). Large numbers of BTS were intentionally introduced from its native range in northern Texas into ponds in the Salinas Valley (Monterey County, California) between 1950 and 1960, where it was subsequently harvested and sold for fishing bait (28). CTS and BTS readily hybridize, and in the time since this introduction, the range of BTS-CTS hybrids (hereafter “hybrids”) has expanded to include much of the Salinas Valley and adjacent grasslands in a hybrid swarm (Figure 1). Established non-native BTS populations in the two other endangered Distinct Population Segments of CTS in Sonoma and Santa Barbara Counties (29) indicate that hybridization affects the entire CTS range, although it is clearly most severe in the central coast region of Monterey County (30).

**Figure 1:**
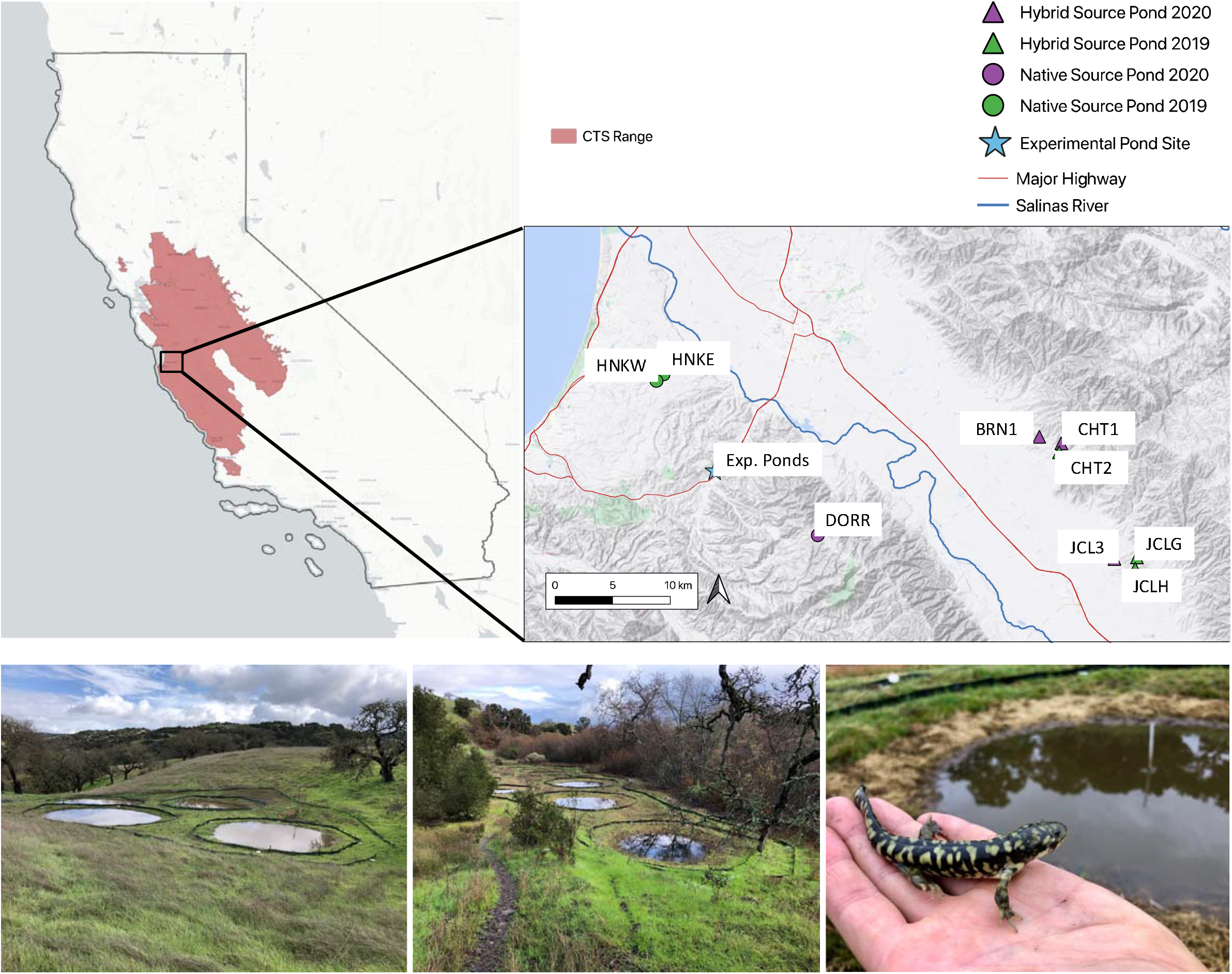
Map of California (on left) with the California tiger salamander range highlighted in red. Map insert (on right) shows an expanded view of the Salinas Valley (Monterey County, CA) with points indicating the location of larval source ponds and experimental ponds. Triangles represent hybrid source ponds and circles represent native source ponds. Green points denote larvae that were collected in year 2019 and purple were from 2020. The blue star indicates the location of the constructed ponds used in the hydroperiod experiment. These ponds are located on the southern edge of the Fort Ord National Monument (Monterey County, CA). Images below depict the naturalistic experimental ponds and a recently metamorphosed hybrid salamander.

Several studies suggest that hybrids enjoy superior fitness compared to pure native CTS. Fitzpatrick et al. (30) modeled the dynamics of hybrid zone expansion and found that non-native alleles reach fixation much faster than is expected based on neutral diffusion models. Other studies have documented hybrid superiority in thermal tolerance (31), locomotor performance (32), and water-quality tolerance (33). This evidence predicts that hybrid salamanders will eventually replace native CTS at least regionally, resulting in genome-level extinction (34). Supporting this view, hybrids are not ecologically equivalent to native CTS, causing trophic cascades that disrupt vernal pool communities (25) and shift the abundance of other native taxa (24) including endangered prey (23). This body of work has emphasized the need for management strategies that reduce or eliminate invasive hybrids in CTS populations.

Effective hybrid management is complicated by the life history of these species. Given the negative effects of hybrids on the landscape, eradication would appear to be a reasonable solution. However, CTS and their hybrids are reclusive, spending the majority of their adult lives in underground rodent burrows, emerging to breed only an average of two times in their life (35–37). This reclusiveness coupled with their 10- to 12-year life span, extreme phenotypic similarity of hybrid and native animals in the field, and average migration distances of 500+ m (with 5% of the population moving 1.8 km) (38), makes eradication and repatriation with native CTS virtually impossible without additional, likely non-traditional, management strategies.

Restoring natural vernal pool functionality has been suggested as a strategy to decrease hybrid advantage. Fitzpatrick and Shaffer (30) found a strong positive correlation between non-native allele frequencies and hybrids breeding in cattle stock ponds with unnaturally long hydroperiods (in many areas, these are their primary breeding sites), and hypothesized that hybrids disproportionately benefit from artificially long hydroperiods. Johnson et al. (39) tested this hypothesis in artificial cattle tanks (mesocosms) by raising different genotypes under varying hydroperiods and found that hybrids enjoyed greater survival and mass at metamorphosis in long-duration mesocosms. Native CTS appeared to fare better than hybrids in the short duration ponds, although this result was less dramatic. Although these results were promising, the experiments were conducted in a highly artificial environment without the normal complement of vernal pool prey and competitive interactions that define this ecosystem.

Here we used large-scale pond manipulations to evaluate habitat restoration as a management alternative to eradication. We tested the effect of variable hydroperiod duration on non-native tiger salamander success under controlled, quasi-natural field conditions. We constructed an experimental array of large seasonal ponds situated at the edge of the hybrid zone to test the hypothesis that shorter hydroperiod ponds favor more native genotypes *in situ*. We inoculated each pond with controlled proportions of early-stage native and hybrid larvae, and evaluated their relative success as they completed metamorphosis and emerged from the ponds. We simultaneously collected genomic data to quantify non-native ancestry, assign survivors to source populations, and interrogate the genome for loci under hydroperiod-mediated selection. This design enables us to evaluate the benefits of managing pond hydroperiod in the field as a potential strategy to minimize the success of non-native genotypes across the landscape.

## Materials and Methods

### Pond Construction

We constructed 18 semi-natural ponds on the Fort Ord National Monument in Monterey County (CA, USA) during September and October 2017, with 14 serving as experimental ponds and four as non-experimental water reservoirs. Ponds filled with rainfall, and we modeled a combination of constructed pond basin and catchment size to produce a gradient of hydroperiods. As in nature, ponds gradually dried down from evaporation. We used a large pump to make minor adjustments to pond volume by adding (from the reservoir ponds) or removing water to achieve hydroperiods from 80 to 115 days. Ponds filled with rain in November, larvae hatched in February and ponds dried from late April – early June in both years (See SI). We installed drift fencing with pitfall traps around each pond to collect post-metamorphic salamanders as they emerged from ponds (See SI; 68, 69), as well as a second, concentric fence array to intercept any trespasses that may have escaped the first fence line. After ponds filled, we inoculated them with plankton from nearby natural ponds in November 2017. After this initial inoculation, ponds were naturally colonized by other vertebrate (e.g. Chorus frogs, *Pseudacris regilla*) and invertebrate salamander prey.

### Experimental Design

Our experimental design consisted of two factors, pond hydroperiod and the relative abundance of non-native larvae. Pond hydroperiod reflects the number of days from larval hatching to pond drying, and represents the total time that the larvae have to develop and complete metamorphosis. Each year there were 7 hydroperiod levels, each replicated twice (14 total experimental ponds per year) each separated by 5 day drying intervals. In year 1 this included ponds that dried in 85, 90, 95, 100, 105, 110, 115 days. Year 2 was accelerated by five days to 80, 85, 90, 95, 100, 105, 110 days (see SI). This range of hydroperiods was based on earlier experimental data demonstrating a significant shift in native/hybrid larval fitness between 90 and 120 days (39). The second experimental factor consisted of the proportions of hybrid to native larvae introduced into each pond: low-hybrid (60 natives:60 hybrids), medium-hybrid (15 natives:60 hybrids) and all-hybrid (0 natives:120 hybrids). Year 1 included one low- and one high-hybrid treatment for each of the 7 hydroperiods, while year 2 included two medium-hybrid treatments per hydroperiod (See SI). These proportions were chosen to span the continuum of larval proportions that exist throughout the hybrid zone. Although differences in these proportions may constitute interesting biological interactions, our study lacks the replication necessary to fully explore such effects, and we therefore include these proportions as a nested effect in our models. We did not include a native-only control treatment for two reasons. First there was exceptionally low levels of native breeding in nearby wild ponds in both years, limiting the number of native larvae we were permitted to remove by the USFWS for this federally and state protected species. Second, previous studies have already explored the optimal conditions for purely native populations (e.g. 23, 36, 40, 69), and our study was focused on the dynamics that occur when hybrids dominate or are actively competing with native genotypes under different hydroperiod regimes. In doing so, our experimental conditions represent the conservation scenarios where the greatest opportunity exists to reverse selection for non-native genotypes.

### CTS and Hybrid Larvae

We collected newly hatched, ~ 15 mm snout-vent length (SVL) pure CTS and hybrid larvae from source ponds around the Salinas Valley. Source ponds were selected based on observed larval abundance and previously-estimated non-native allele frequencies (43). We selected five ponds in 2019 and five different ponds in year 2020 for a total of 10 unique source ponds in the study (Figure 1), all within ~ 30 km of our experimental ponds. After capture, larvae were immediately sorted into larger and smaller size classes, transported to the experimental ponds in their natural pond water, and allowed to acclimate to the experimental pond conditions for 1 hour to minimize transition shock. In each year, the number of larvae of each size class from each source pond was balanced across treatment ponds to maximize the probability that each experimental pond started with equivalent allele frequencies. The larval densities used in this experiment are within the range of natural CTS densities, which a previous study estimated to be between 3.5 and 7.0 larvae/m^3^ (See SI; 25). We also retained a representative sample of 40-60 larvae, comprised of the same proportion of large and small size classes, from each source pond, which were immediately euthanized and later sequenced and used as reference genotypes to assign experimental animals to source ponds.

Larvae developed through metamorphosis and naturally emigrated from ponds, where they were intercepted by drift fencing surrounding each pond and captured in pitfall traps. Open traps were checked each morning before sunrise. At the time of capture, we collected 1) time and date; 2) bucket location; 3) metamorph total length (mm); 4) snout-to-vent length (mm); 5) mass (g); and 6) a sample of genetic tissue (1 cm of tail tip). Length was measured to the nearest millimeter using a standard ruler, and mass using a digital scale (0.01g precision). All metamorphs were euthanized using a 5g/L solution of tricaine methanesulfonate (“MS-222”; (44) and preserved in the UCLA HBS museum.

### Molecular Methods and Bioinformatics

We extracted genomic DNA from reference source pond larval and experimental metamorph tissue using a modified salt extraction protocol (45). DNA was diluted to 100 ng/uL (10,000 ng total) and sheared to approximately 500 bp using a BioRupter (Diagenode, Denville, NJ). We performed a double-sided size selection using SPRI beads (46) to obtain an average fragment size of 400 bp, and recovered approximately 1,000ng of DNA to use in library preparations. We used Kapa LTP library preparation kits (Kapa Biosystems, Wilmington, MA) to perform standard Illumina library preparations (end repair, A-tailing, and adapter ligation). Sample libraries were dual-indexed using 8-bp indices that were incorporated using PCR (adapters from Travis Glenn, University of Georgia), and combined into pools (16 individuals, 250 ng/individual, 4,000 ng total DNA in 7uL in 10mM Tris-HCl, pH 8) for sequence capture reactions targeting 5,237 loci with a CTS-specific protocol (47). Each pool was sequenced on an Illumina NovaSeq S4 150-bp paired-end lane at the Vincent J. Coates Genomics Sequencing Laboratory at UC Berkeley. Adapter sequences and low quality bases were trimmed from raw sequences using trimmomatic (48). We used bwa-mem to map trimmed reads to the published Mexican axolotl (*Ambystoma mexicanum*) genome (49, 50), a close relative of CTS and BTS (51, 52). We then followed the gatk (version 4.0) best practices pipeline (53) for calling variants across all samples (See SI).

We calculated the Hybrid Index Score (“HIS”; 54) for each source pond sample and each surviving metamorph. To calculate the HIS we used a reference panel comprised of 150 confirmed native CTS that span the species’ range and 30 non-native BTS individuals from the original source population that was introduced to California (30). We identified diagnostic loci that are fixed (or essentially fixed)-different between the two species following established protocols (See SI; 31). For each individual, we then calculated the HIS as the proportion of BTS-derived alleles divided by the total number of non-missing alleles scored in that individual. For each experimental pond, we estimated initial HIS by taking the average HIS of each source pond weighted by the number of larvae added from each source pond.

### Differential Group Survival

To test for the effect of larval source pond on survival, we assigned metamorphs to their original source pond using Discriminant Analysis of Principal Component (DAPC). For each year of the experiment we used the representative sample of source pond larvae to construct a DAPC using adegenet in R version 4.0.4 (55). This model determines the principal component axes (eigenvectors) of genetic variation that best discriminate between the pre-assigned source populations. If two or more populations were not easily distinguishable in the first DAPC, we ran additional subset DAPC models with only the overlapping populations and metamorph genotypes. We used these DAPC models to predict the pond of origin of all metamorphs that emerged from the experimental ponds. Six individuals that could not be assigned to a single source pond were dropped from the analysis.

We identified likely sibling cohorts (“family groups”) from each source pond and metamorph groups using the program Colony 2 (56). We ran Colony separately for each source pond, comprised of source pond larvae and the metamorphs previously assigned to that source pond. We analyzed the “BestCluster” output files from Colony to determine probable family groups within each pond group using custom R scripts and used these cohort assignments for downstream analyses.

We investigated the possibility that experimental pond hydroperiod drives differential survival among source ponds or family groups. For each experimental pond we calculated a chi-squared statistic (*χ*^2^) to quantify the degree of dissimilarity between the input (pre-selection) and output (post-selection) group distribution. We performed this analysis for source pond and family groups. For each, we used the known input proportions as expected values, and calculated observed proportions in the metamorphs from our computational assignments to either family group or source pond. We used Linear Mixed Model (LMM) regression using the lmer package in R to test whether larval survival through metamorphosis was associated with source pond or family, modelling the log-transformed *χ*^2^ statistic as the response variable with hydroperiod as the predictor and larval proportion as a nested random effect.

### Larval Proportions

The different proportion of native to non-native larvae was included in each analysis as a nested random effect in our hierarchical models. We did this because the focus of our experiment was to explore the fitness effects of hydroperiod and individual genotype that exist in ponds throughout the natural hybrid swarm, which include a wide range of native and non-native larval proportions. We therefore consider the larval proportion to represent draws from the natural variation in pond composition. Including larval proportion as a random effect accounts for these nested differences, enabling us to compare the main effects of hydroperiod and HIS at the level of the individual.

### Simulating Input Larval Genotype Composition

Analysis of genotype-specific larval survival requires an accurate estimate of the composition of larval genotypes that were added to each experimental pond (“input larvae”), and we were not able to take tissues from the actual larvae that were added (they are too small and delicate to non-destructively sample). Instead, we reconstructed the expected genotypes of input larvae based on the genotyped a sample of 40-60 larvae from each representative source pond. We know the exact proportions of larvae added from each source pond, and can estimate the family group composition of each source pond based on our representative sample. We randomly sampled genotypes from the predicted sib-groups, while maintaining their relative proportions, to accurately simulate the allelic composition of the larvae that went into each experimental pond, ensuring that overall allele frequencies are maintained. We then combined these simulated data with the true survivor data (metamorphs) to produce a full, genomically-informed, before-and-after selection dataset for use in the survival analysis. We repeated this process 500 times with different random draws from the representative larval group, performed an independent Bayesian analysis on each iteration, and combined the resulting posterior distributions to account for the unknown variation that may arise from different genotypes of input larvae.

### Larval Survival

We modeled factors predicting larval survival using a hierarchical Bayesian framework. Our likelihood function included pond hydroperiod (HYDP) and larval HIS (HIS) to predict the probability of survival (p). An interaction term between HYDP and HIS was initially included, but removed if it was non-significant and reduced model fit. Predictors HYDP and HIS were both centered and scaled using the scale function in R. The larval proportion group (t) was included as a random effect, such that the intercept b_0_ was determined for each group separately. See supplemental for additional Bayesian model specifications (See SI).

### Metamorph Mass

We employed a hierarchical Bayesian model to explore the effect of Hydroperiod and HIS on metamorph mass. Our likelihood function included a quadratic term, hydroperiod-squared (HYDP^2^), hydroperiod (HYDP) and larval HIS (HIS) to predict metamorph mass (MASS). An interaction term between HYDP and HIS was initially included, but removed if it was nonsignificant and reduced model fit. The predictors, HYDP^2^, HYDP, and HIS were centered and scaled using the scale function in R. The larval proportion group (t) was included as a random effect, such that the intercept b_0_ was estimated for each group separately. See supplemental for additional model information (See SI).

### Standardized Mass

We also analyzed differences in mass at the pond-level using a hierarchical Bayesian model. We standardized the metamorph mass based on the number of larvae that were initially added to the pond, to account for differences in larval density in 2019 and 2020. This standardized value (Mass per Input Larva; “MPIL”) facilitates the comparison of overall metamorph biomass while controlling for initial larval stocking densities. We removed zeros in MPIL by adding half of the lowest recorded MPIL to every MPIL value in the dataset and then log-transformed the values. We split larvae into two discrete genotype categories: native (HIS < 0.10) and hybrid (HIS ≥ 0.10). Our likelihood function included pond hydroperiod (HYDP) and larval Genotype (GEN) to predict MPIL. An interaction term between HYDP and GEN was initially included, but removed if it was non-significant and reduced model fit. The predictor HYDP was centered and scaled using the scale function in R. The larval proportion group (t) was included as a random effect, such that the intercept b_0_ was determined for each group separately. See supplemental for detailed model specifications (see SI).

### MCMC Implementation

For all Bayesian analyses we used the R package jagsUI (version 1.5.2) which implements a Gibbs Sampler in the R environment. Unless otherwise indicated, all Bayesian models were iterated 10,000 times with four independent chains. We allowed automatic adaptation and specified zero-iteration burn-in. Bayesian model convergence was assessed using the Potential Scale Reduction Factor (PSRF or Rhat 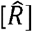), which we required to be less than 1.05. We visually inspected the MCMC output using the R package MCMCoutput (version 0.1.1) to ensure adequate convergence and normal posterior distributions. Parameter significance was determined if the 95% Credible Interval (CI) did not include zero. Parameter estimates were taken as the mean value of the posterior distribution.

### Loci Under Selection

We used the program BayEnv (57) to scan the sequenced target regions for loci that experienced differential selection across the hydroperiod treatments. Since we could not employ hierarchical modeling we analyzed metamorphs from each of the three larval proportion groups separately, using hydroperiod as a continuous predictor. BayEnv calculated a Bayes Factor for each locus after controlling for the underlying variance-covariance structure that results from uneven sample sizes and shared population history (58). These Bayes Factors are used in Bayesian model selection and can be interpreted similarly to the frequentist likelihood ratio (59). Following Jeffery’s scale of evidence for Bayes factors, we selected loci with a Bayes factor of 10 or greater, which suggests “very strong” evidence for selection (See SI; 84). We then analyzed the raw allele frequencies of these significant loci to determine the direction of selection. We built GLMs with the frequency of reference alleles (# reference / (# reference + # alternate)) as the dependent variable and hydroperiod as the independent predictor. We extracted the model coefficients to determine the direction (+/−) and magnitude (absolute value) of the shift in allele frequency. We also used the frequency of reference and alternate alleles in our reference panel of 150 pure CTS and 30 pure non-native BTS to identify alleles that were predominantly associated with CTS or BTS. We tested for differences in the number of loci that experienced a CTS versus BTS biased shift in allele frequencies using a 1-sample proportions test. Finally, we constructed a linear mixed model to test the relative strength of the allele frequency shift in CTS versus BTS associated alleles. This model included the absolute value of the marginal effect of allele frequency shift as the response variable and the genotype of the preferred allele as the predictor, with larval proportion as the random effect.

We investigated the possibility of functional enrichment among genes that experienced hydroperiod-mediated selection using the web server “g:Profiler” (61) to identify gene ontology terms that were overrepresented in the BayEnv list. We compared our list of significant genes to all of the genes in our 4,723 gene panel that were annotated in the closest well-annotated model organism, the African clawed-frog (*Xenopus tropicalis*) in addition to the best studied system, humans (*Homo sapiens*). We report the gene ontology terms that had a p-value of less than 0.05 after applying the software’s custom “g:SCS” correction for multiple tests.

## Results

### Field Results

The experimental ponds successfully held water and achieved the desired range of hydroperiods. In 2019 ponds held water for 85 to 115 days and in 2020 between 80 and 110 days. Across years, 249 living tiger salamander metamorphs or near-metamorphs were recovered (Figure 2). Twenty-one were found in dried pond basins and had failed to complete metamorphosis; these animals were sampled, but excluded from all analyses, since they presumably would have succumbed to heat and desiccation in the wild. The remaining 228 successful metamorphs constitute an across-year survival rate of 8.4% out of the 2,730 larvae that were originally introduced into the experiment. Survival rates were similar in 2019 (149 metamorphs out of the 1680 total larvae; 8.9% survival rate) and 2020 (76 metamorphs out of 1,050 initial larvae; 7.2% survival rate). Although no studies have specifically examined larval-to-metamorph survival rates, these values are reasonable given field-based estimates of egg-to-metamorph survival rate of 2.5% (Searcy et al. *in review*). Across years the median metamorph mass was 9.45g ± 0.29 (standard error), which agrees quite well with long-term field data which estimated average metamorph mass to be 9.4g (Searcy et al. *in review*).

**Figure 2:**
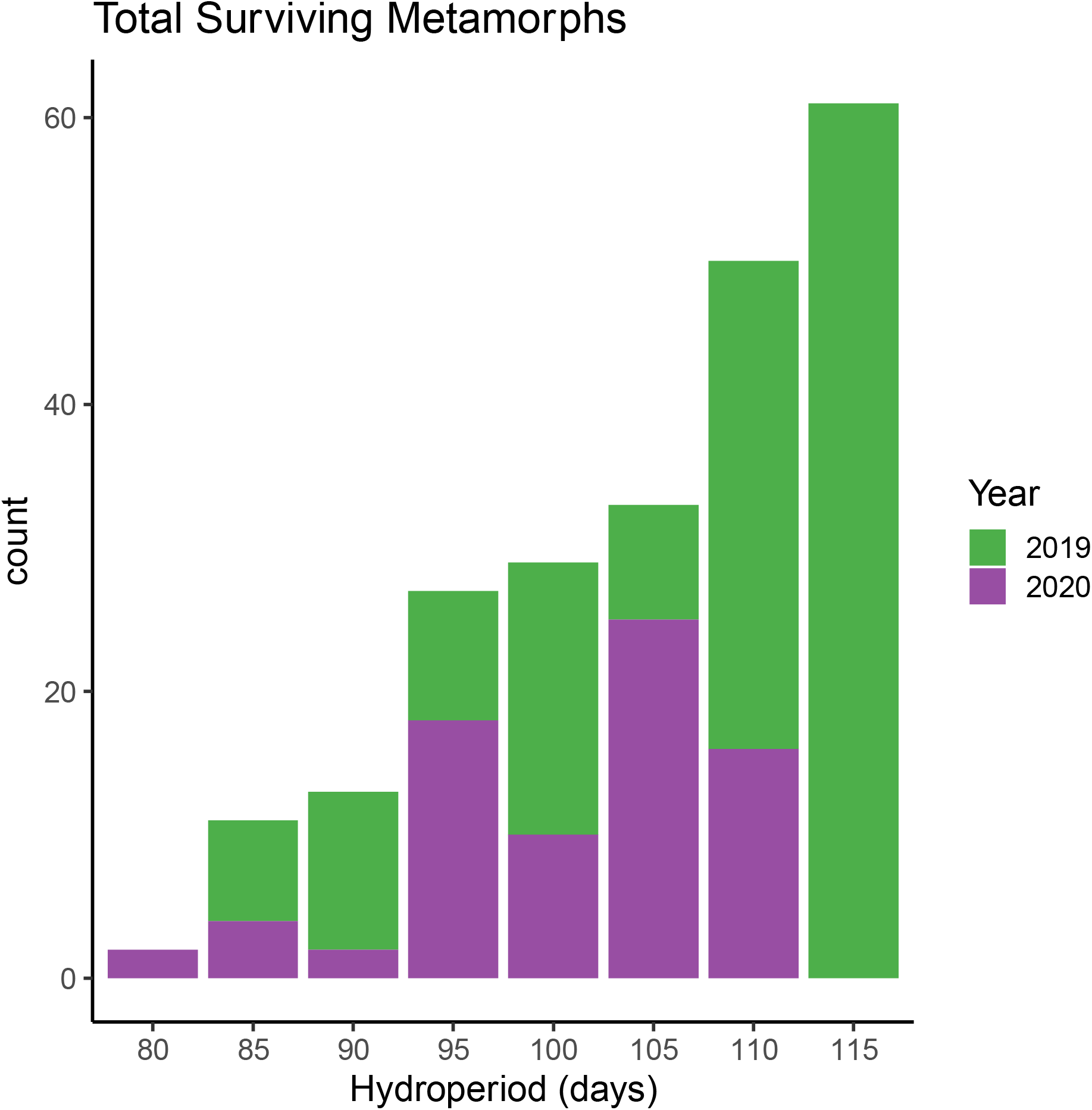
The total number of successful metamorphs captured in 2019 and 2020 across hydroperiod treatments.

### Sequencing Results

We sequenced a total of 485 representative larvae from the 10 source ponds (median = 58/pond-year) and 249 metamorphs for a total of 734 individual salamanders. Of these, 27 source pond larvae and 1 metamorph had low sequencing depth and were dropped. We generated 1,732 billion base pairs of sequence data across 5.74 billion read pairs, identified 255,350 single nucleotide variants (SNPs), which yielded 5,919 SNPs after filtering and 1,788 CTS/BTS diagnostic SNP markers (See SI).

### Source Pond Hybrid Index Score

We identified non-native alleles in all larval source ponds that were expected to contain hybrids based on previous field-based sequencing (40). However, we found that hybrid ponds consistently had a much greater degree of non-native ancestry than expected based on earlier results. Earlier genomic surveys from the last two decades found that these ponds often had HIS between 0.5 and 0.8. However, the average HIS across hybrid source ponds was 0.90 ± 0.065 (median ± SD; Figure 3). While this does not capture the range of HIS that we had anticipated, it does accurately represent a snapshot of the current HIS dynamic in these hybrid zone ponds. The average HIS of native source ponds was 0.07 ± 0.016. This small degree of apparent hybridity is almost certainly due to sample missingness, incomplete lineage sorting compared to the reference panel, or other errors in considering our marker loci “fixed” between species. We therefore consider individuals with HIS < 0.10 to be native and individuals with HIS ≥ 0.10 to be hybrid for all further analyses.

**Figure 3:**
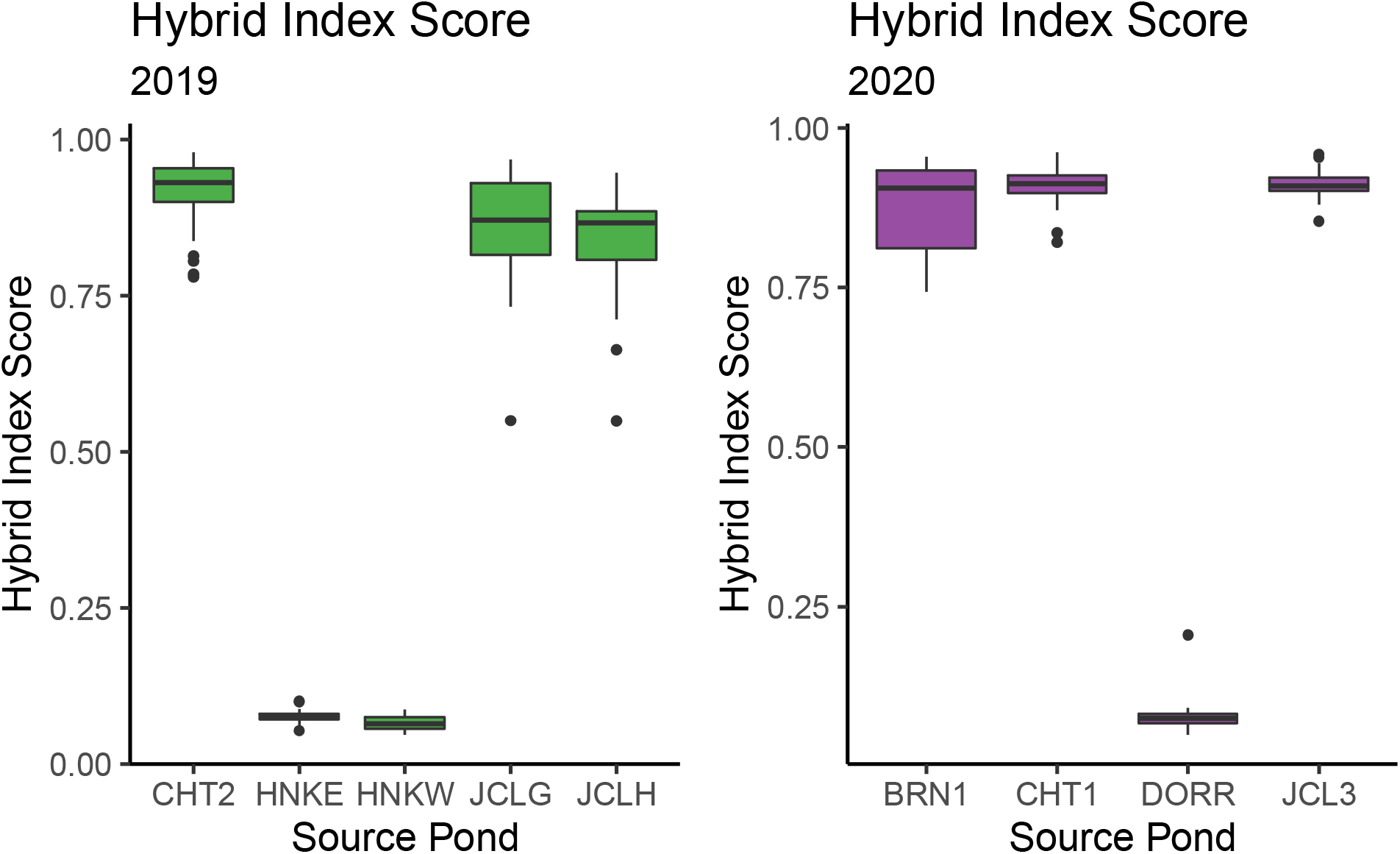
The Hybrid Index Score (“HIS”) of each of the source ponds from which larvae were collected. HIS is scaled from 0 (completely native CTS) to 1 (completely non-native BTS).

### Differential Group Survival

We found a significant increase in source pond dissimilarity (*χ*^2^) as hydroperiod increased (LMM: estimate = 0.03, CI =(0.006, 0.063), p = 0.02, Figure 4A). This increase in dissimilarity indicates that the surviving larvae in longer duration ponds are not evenly distributed across the initial groups that were added to each pond, indicating among-pond-cohort selection that becomes more pronounced in longer duration ponds. We found a similar, but weaker, change in the distribution of individuals across family groups (LMM: estimate = 0.029, CI = (0.002, 0.056), p = 0.047; Figure 4B). A graphical representation of this non-random survivorship can be viewed in the supplemental Figure SI.1.

**Figure 4:**
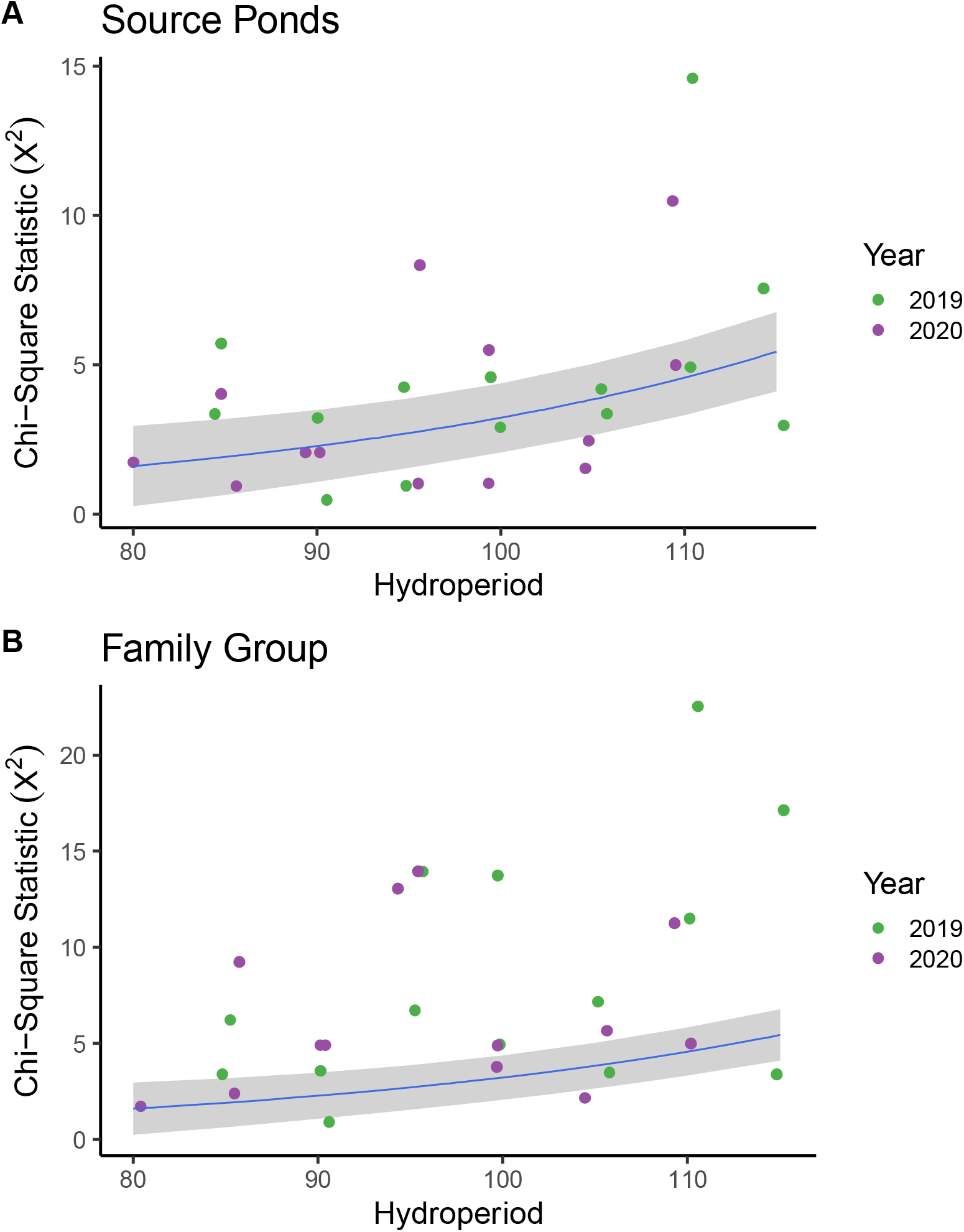
Effect of hydroperiod on surviving source pond and family group distributions. The response is *γ*^2^ which represents the departure from the starting larval group proportions, calculated as the sum of (observed-expected)^2^ / expected). Points were jittered on the x-axis to show overlapping values. The panels show the increase in source pond (A) and family group (B) dissimilarity as pond hydroperiod increases. These results suggest that the larvae that survive in longer duration ponds are not evenly distributed across the groups that were added to each pond. This non-random distribution of survivors may indicate kin-level selection that becomes more pronounced in long duration ponds. Blue line represents the log-linear regression line with grey ribbon depicting standard error.

### Larval Survival

Larval survival significantly increased with longer pond hydroperiod (Bayes: b_HYDP_ = 0.08, CI= (0.06, 0.10), Rhat = 1.0) and with greater HIS (Bayes: b_HIS_ = 0.53, CI= (0.006, 1.05), Rhat = 1.0). The interaction term between hydroperiod and HIS was not significant (b_INT_ = −0.04, CI = (−0.23, 0.139)) and decreased model fit, and was therefore removed. These results indicate that hybrids maintain a survival advantage over native CTS across all levels of hydroperiod included in this study. Based on our model predictions, survival for a 90% non-native hybrid (HIS = 0.90) increased from 2.1% in an 85-day hydroperiod, to 23.7% survival in 115 days, a roughly eleven-fold increase (Figure 5A). The effect of this increase is biologically significant, given the average larval survival rate of 8.4% in this study. Native CTS survival was lower overall, but followed a similar trend with respect to hydroperiod, increasing from about 1.3% survival at an 85-day hydroperiod to 14.7% at 115 days, also an eleven-fold increase (Figure 5B).

**Figure 5:**
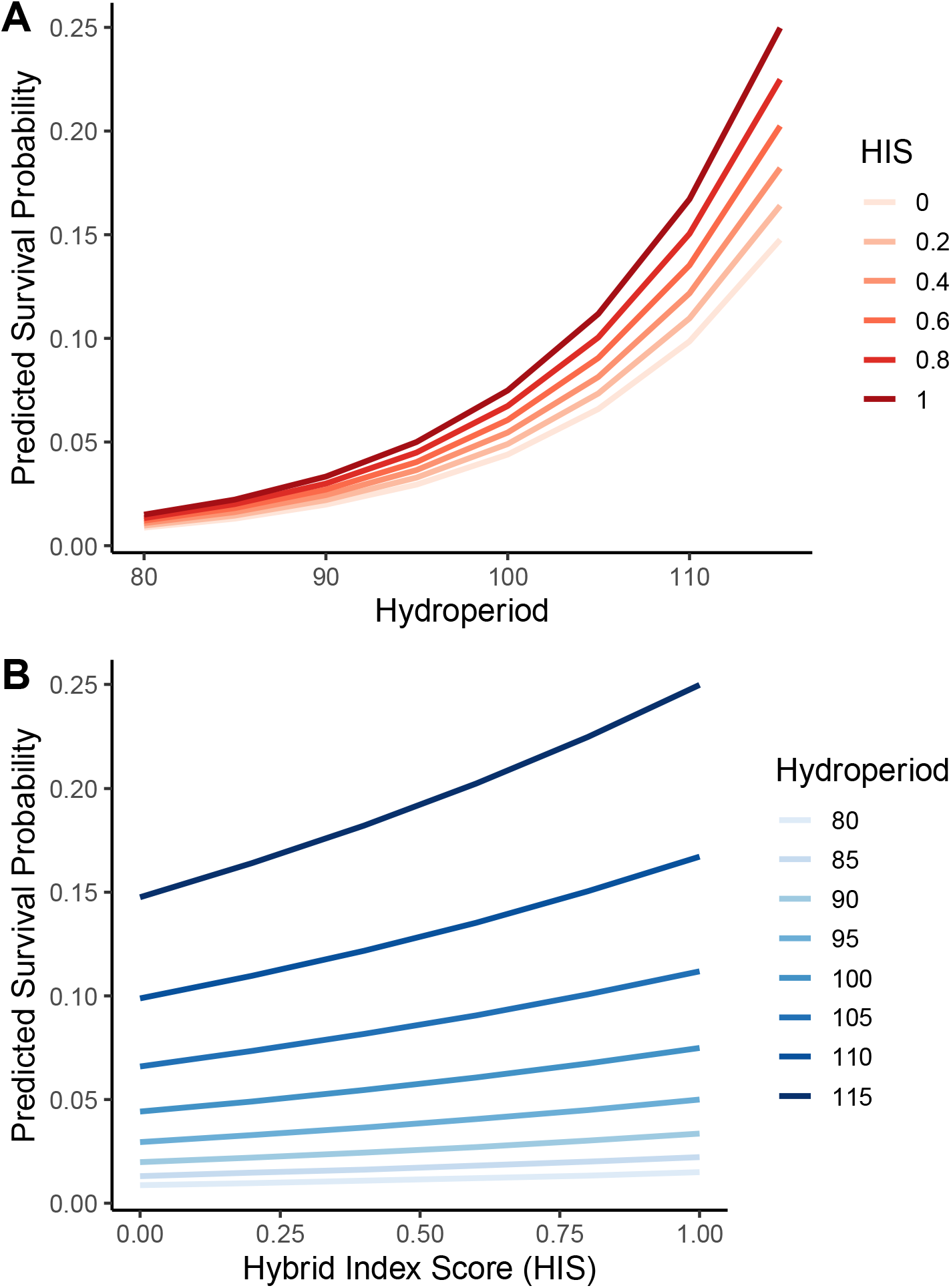
Model predictions for larval survival probability based on pond hydroperiod and hybrid index score (“HIS”). Representative samples of all wild source pond larvae were resampled using genomic data to reconstruct the complete assemblage of larvae that were added to the experimental ponds. Genomic analyses ensured an accurate representation of input larvae down to the sibling-group level. We iterated this process 500 times to capture all potential variation inherent in the resampling procedure. We constructed a hierarchical Bayesian model for each iteration and combined the posterior distributions for each model parameter. Panels show the model predictions across a range of hydroperiod (A) and HIS (B).

### Metamorph Size

Mass at metamorphosis was significantly correlated with HIS (Bayes: b_HIS_ = 2.37, CI= (1.88, 2.87), Rhat = 1.0; Figure 6B), hydroperiod (Bayes: b_HYDP_ = 12.25, CI= (3.43, 21.26), Rhat = 1.0; Figure 6A) and hydroperiod^2^ (Bayes: b_HYDP_^2^ = −12.21, CI= (−21.08, −3.19), Rhat = 1.0; Figure 6A), which was included in the model to account for non-linearity in mass. The interaction between hydroperiod and HIS was not significant (b_INT_ = −0.37, CI = (−0.83, 0.08)) and reduced model fit, and was therefore removed. We show the relationship between metamorph mass and HIS (Figure 6B) and metamorph mass and hydroperiod + hydroperiod^2^ (Figure 6A) separately to improve interpretability. The predicted mass of a native CTS (HIS = 0.05) at metamorphosis was 3.5g at an 85-day hydroperiod, which increases to 4.7g at 115 days, with a mean of 4.6g across treatments. The predicted mass of a predominantly non-native hybrid (HIS = 0.90) was 10.1g at 85 days which increased to 11.3g at a 115-day hydroperiod, with a mean of 11.1g across treatments.

**Figure 6:**
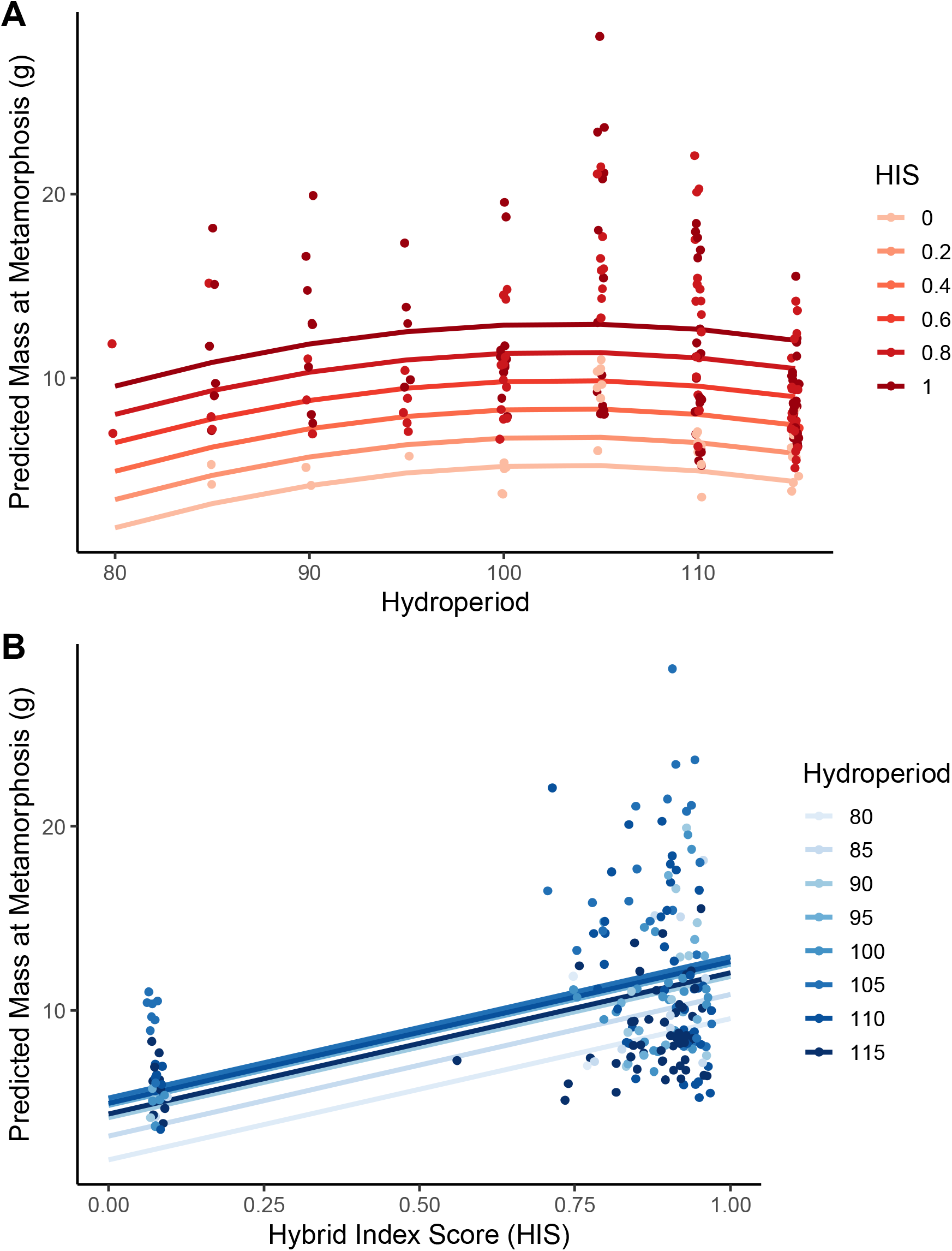
Model prediction for the effect of hydroperiod and hybrid index score (HIS) on metamorph mass. We used a hierarchical Bayesian model which incorporated HIS, hydroperiod and hydroperiod^2^. The quadratic term was included to account for the non-linearity in mass as hydroperiod increases.

There was a significant increase in the standardized metamorph mass with respect to hydroperiod (Bayes: b_HYDP_ = 0.52, CI= (0.26, 0.78), Rhat = 1.0; Figure 7), and genotype (Bayes: b_GEN_ = −1.41, CI= (−1.95, −0.88), Rhat = 1.0; Figure 7). The interaction between hydroperiod and genotype was not significant (b_INT_ = 0.02, CI = (−0.51, 0.55)) and reduced model fit, and was therefore removed. Based on model predictions, hybrid ponds with an 85-day hydroperiod were predicted to produce 0.33g of metamorph per input larva (“MPIL”; g/input larva) which increases to 1.89g/input larva at 115 days. In contrast native ponds were only predicted to yield 0.02g/input larva at an 85-day hydroperiod, rising to 0.40g/input larva at 115 days. To put these values into ecological context, we can calculate the total predicted output of a female by multiplying by 814, the average clutch size for native CTS (41) and by the estimated egg survival rate of 78% (unpublished data), resulting in roughly 635 viable larvae per female breeding event. This means that a single hybrid female would produce 210g of metamorph in an 85-day pond or 1,200g in a 115-day pond. In contrast, a native CTS female would produce 13g in an 85-day pond and 254g in a 115-day pond.

**Figure 7:**
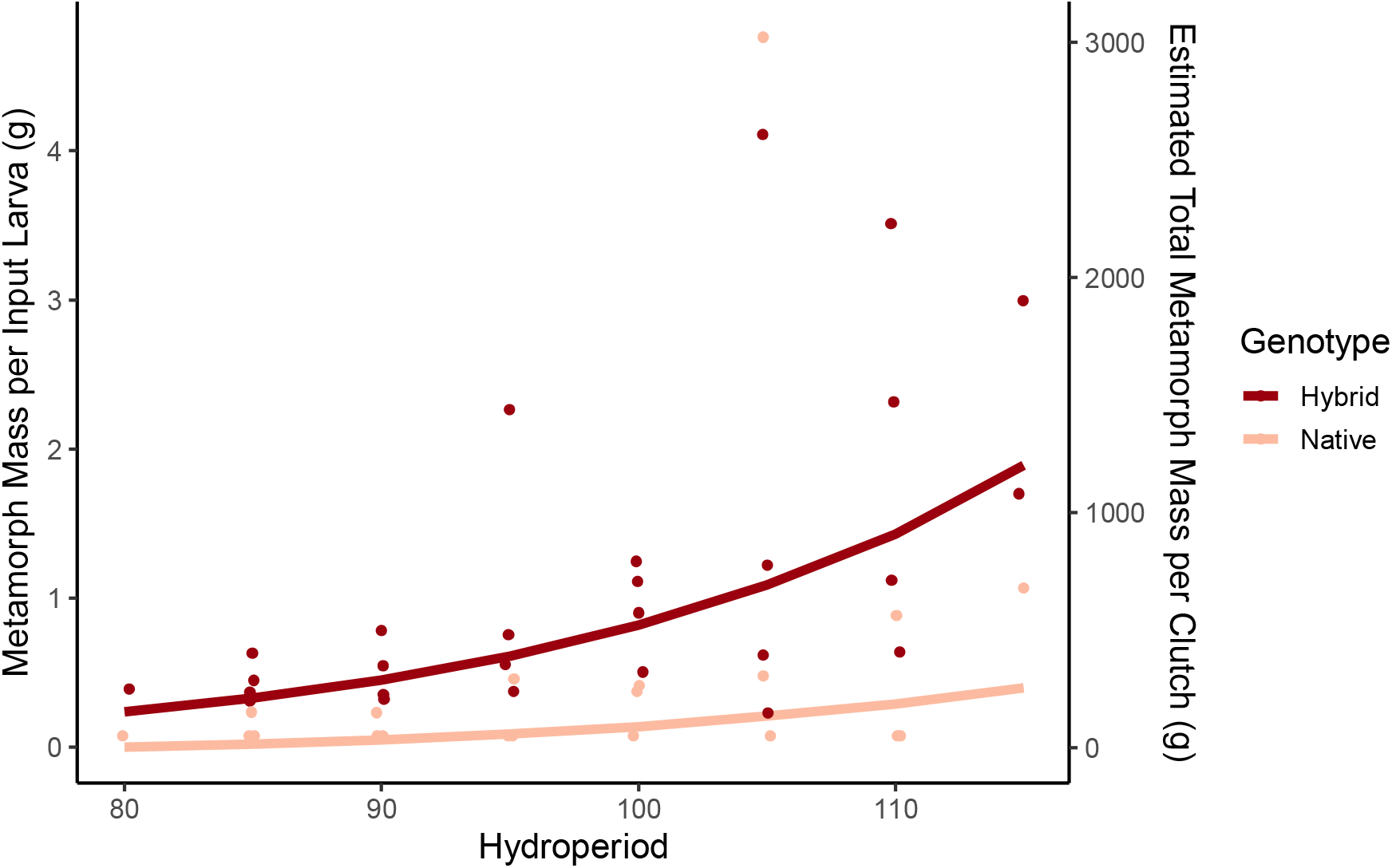
The relationship between overall metamorph mass produced from a pond and hydroperiod. Metamorph mass per input larva (MPIL) standardizes the overall metamorph biomass to facilitate comparisons across years. This value is calculated as the sum of all metamorph mass that emerged from a pond, divided by the number of larvae that were added to the pond. This standardized metric enables us to compare overall productivity across treatment types. To put this value into ecological context, on the right axis MPIL is multiplied by the average number of larvae produced per female clutch (635, see Methods). This value represents the estimated total grams of metamorph each female would produce given their genotype and pond hydroperiod. Each point represents the standardized mass of a specific genotype for a given pond. Points are colored by genotype, where HIS < 0.1 is native (light red) and HIS ≥ 0.1 is hybrid (dark red).

### Loci Under Selection

We identified 86 loci that exhibited significant allele frequency shifts resulting from the hydroperiod treatment, including 22 loci (9 BTS and 13 CTS biased shifts in allele frequency) in the “low-hybrid”, 34 loci (12 BTS and 22 CTS) in the “medium-hybrid”, and 30 loci (17 BTS and 13 CTS) in the “all-hybrid” groups with a Bayes factor greater than 10. For context, these 86 loci represent 1.8% of the 4,723 annotated genes included in our study. We observed a single gene (“SRFBP1”) that was significant in more than one larval group. There was no difference in the number of loci that experienced CTS or BTS biased shifts in allele frequency (Prop Test: p = 0.33, prop = 0.56, CI = (0.45, 0.66)). However, loci that exhibited a CTS biased shift had a greater magnitude of effect (i.e. greater increase in CTS allele frequency) than BTS biased loci (LMM: estimate = 3.40×10^-3^, CI = (1.4×10^-3^, 5.2×10^-3^), p = 5.1×10^-4^). There were no enriched gene ontology (“GO”) terms in our list of loci under selection when compared with the *Xenopus tropicalis* annotation library. There was, however, a single significant GO term (“hsa-mir-10b-5p”) when compared to the human annotation repository (corrected p = 3.71×10^-2^).

## Discussion

Extinction via hybridization is both real in nature and controversial in terms of its perceived importance to conservation (62). Regardless of one’s views on the conservation threat of hybridization (21, 63), it is clear that once hybrids are established, they can be difficult or impossible to eliminate, given that their ecological and physiological impacts reside at the genic, rather than whole-organism, level. To even quantify the presence of hybridization often requires sophisticated genomic tools. Estimating its impact often requires field-based ecological experiments to evaluate fitness, both in absolute terms and compared to the native parental lineages that are being replaced (21, 25, 31, 33). If hybrids are viewed as a conservation threat, as has been the case in many systems, the most vexing challenge is the final step—how to differentially reduce or eliminate non-native alleles from an admixed hybrid swarm, or at least slow the advance of non-native introgression in the wild. Our whole-pond experiments took on this challenge, building on observational field data suggesting that extended hydroperiods, which are the hallmark of human-modified salamander breeding sites, are the causal reason for hybrid success in central California. Our results were mixed: in experimental treatments that mimic those at the advancing front of the hybrid swarm, we confirm that 1) across hydroperiods, hybrids always outperform native salamanders, and 2) longer hydroperiods produce more fit metamorphs regardless of genotype. Over the short-medium hydroperiods that we tested, we found no evidence that hybrids were disproportionately favored in longer duration ponds. Genomic data comparing surviving metamorphs to the input gene pool identified candidate genes that may underlie the response to extreme hydroperiod pressure in natives and hybrids alike. Although not a silver bullet for reducing or eliminating non-native genotypes, our experiments point to pond hydroperiod as a powerful tool to enhance or decrease salamander fitness for both native and hybrid genotypes.

### Differential Group Survival

Across hydroperiods, differential survival shifted the distribution of successful individuals from the initial proportions of larval source pond and family groups (Figure 4). It appears that only a few source ponds/family groups dominate the experimental pond survivors in longer-duration treatments (See SI: Figure SI.1). This could result from a number of different mechanisms, including kin recognition and family-level selection. Pfennig et al. (64) showed that cannibalistic morphs of BTS preferentially consumed unrelated larvae, and later confirmed kin selection as the mechanism (65). Although the exact mechanism of this kin selection appears variable during different life stages (66), it is consistent with the reduction in source pond and family diversity that we observed as hydroperiod increases. Recent work has demonstrated that food abundance in natural ponds decreases in the late larval season, and that late-stage larval CTS exclusively consume the largest available prey irrespective of whether they are invertebrate or vertebrate taxa (23). This would likely increase rates of cannibalism, which has rarely been documented in CTS, but is common in other members of the tiger salamander complex, including BTS (23, 24, 67). If selective cannibalism that avoids close kin occurs in long hydroperiod ponds with a mix of hybrid and native CTS, it could create a positive feedback loop where native CTS experience progressively greater predation as the number of hybrids in a population grows. This dynamic may be exacerbated by the substantial size difference we detected between native and hybrid larvae, facilitating cannibalism of the smaller natives. Future research aimed at experimentally determining cannibalistic interactions between hybrid and native CTS, and their relation to pond hydroperiod, should be a high priority, and may help explain the rapid turnover of ponds from native to hybrid genotypes; it may also indicate a previously unrecognized cost to long hydroperiod ponds in the hybrid swarm.

### Larval Survival

Hybrids enjoy a greater probability of survival across all hydroperiods examined here, corroborating previous studies. Fitzpatrick & Shaffer (68) found that increased heterozygosity enhanced the survival of hybrid larvae in wild populations, which may explain this consistent superiority. While the degree of enhanced hybrid survival in long hydroperiod treatments agrees with previous mesocosm-based experiments (39), we found no indication that hydroperiod affected native vs. non-native survival differently. Instead, we show that both genotypes enjoy the same relative benefit from increased hydroperiod.

There are several potential explanations for this apparent lack of concordance with earlier results. First, our experiments were conducted under essentially natural conditions, including natural predation, competition, prey abundance, and abiotic influences. It may be that native CTS have greater survival in short duration ponds only under artificial conditions, and that this effect disappears in the face of strong competition with hybrids. If so, this underscores the importance of conducting field-based ecological experiments that accurately replicate the conditions experienced in the wild, especially for applied conservation research. Second, previous mesocosm experiments included only a single genotype per treatment (e.g. all CTS, all BTS, all hybrid), which does not accurately reflect pond compositions throughout the hybrid zone, specifically at the leading edge where few hybrid migrants colonize pure native ponds. Our study examines the fitness of native CTS in the context of mixed-genotype ponds which represent the most pressing management scenarios. Third, although the previous mesocosm study did report greater survival for natives vs hybrids in short hydroperiods, it also reported very low native survival in the longest hydroperiod treatment (greater than 150 days, approximately 30% survival) compared to the short treatment (90 days, approximately 85% survival), a perplexing result. It is possible that this large effect drove the significant interaction term between genotype and hydroperiod. We did not include such a long hydroperiod treatment (our longest was 115 days), because our study was designed to explore the most potentially useful conservation outcome; identifying a short hydroperiod treatment where native CTS outperform hybrids. Thus, it may still be true that hybrids benefit disproportionately from hydroperiods longer than 115 days, in which case reducing such hydroperiods could remove some of the additional benefit hybrids enjoy. This would be consistent with field-based observational studies documenting greater non-native allele frequencies in artificially deep, perennial ponds (30, 69). If true, eliminating very long hydroperiods would not reverse selection for non-native genotypes, but it would slow the spread of non-native alleles, buying time for other management actions to be initiated. Future studies should explore this potential management strategy.

### Metamorph Size

Size at metamorphosis is the most important single predictor of survival and fitness in ambystomatid salamanders generally, and in CTS specifically (42, 70). Metamorph mass and length were both positively correlated with HIS and hydroperiod (Figure 6B), demonstrating that greater proportions of non-native ancestry produce larger metamorphs. Averaging across hydroperiod treatments, our analyses predict that hybrids will be 2.4x more massive than native CTS over the same larval life-span and environmental conditions. This disparity in mass likely plays a tremendous role in the apparent fitness advantage of adult hybrids in the wild. Previous work has highlighted the critical role of mass at metamorphosis in the survival and lifetime fitness of native CTS (42), and assuming that this also applies to non-native hybrids, this genotype-linked size difference may well be driving selection for non-native genotypes in the field.

Mass at metamorphosis was also correlated with pond hydroperiod, though not in a simple monotonic relationship (Figure 6A). The quadratic hydroperiod term in the model accounted for the apparent drop in metamorph mass in the longest hydroperiod ponds (Figure 6A). It is likely that this modest decrease was driven by 1) the increase in the number of metamorphs that survived under longer hydroperiod conditions, combined with 2) a decrease in prey abundance towards the end of the season driven by a combination of normal seasonal declines in some taxa (23) and 3) elevated rates of within-pond competition for prey. Previous work on a related congener, *Ambystoma talpoideum*, similarly found that increased pond duration did not have a consistent effect on individual mass at metamorphosis (71, 72). Instead, longer hydroperiod ponds produced a greater number of individuals that successfully completed metamorphosis.

When we compared total metamorph mass that emerged from each pond after correcting for the number of larvae that were initially added, we found that longer hydroperiods produced more metamorph biomass overall and that ponds with hybrid salamanders produce substantially more offspring biomass than native ponds over the same duration. At an 85-day hydroperiod, hybrid females produce roughly 16.5x more offspring biomass than native CTS females. This advantage decreases to 4.7x in the longer 115-day hydroperiods ponds. It appears that the enormous disparity in offspring production in the short duration ponds stems from the low expected offspring mass for native CTS at just 0.02g/input larvae or 13g of predicted offspring per female. Based on previous mesocosm studies (39), we predicted that native CTS would have an advantage in the shorter duration ponds since they have evolved in the California landscape, adapting for millions of years to a Mediterranean climate characterized by sparse rain, elevated temperatures, and naturally short pond duration (31). Many factors may explain this disparity, including competition or cannibalism when CTS and hybrids develop in the same pond, as was always the case for pure CTS in our experiments. Other potentially important factors include unrecognized importance of long-hydroperiod playa lakes in the long term stability of native CTS metapopulations (40), or the possibility that, through introgression and selection, hybrids have already assimilated the alleles that underly short pond adaptation. Coupled with the apparent increase in fitness from elevated heterozygosity inherent in hybrids (68), this may have produced the transgressive ‘hopeful monsters’ (73) that exceed the fitness of native CTS, as has been demonstrated in thermal tolerance (31).

### Loci Under Selection

Given the clear effect of hydroperiod treatments on fitness, we interrogated our exon panel for candidate genes that may be responding to selection from pond duration. We identified 86 candidate genes that may have undergone selection in response to hydroperiod through the course of our experiment. Although exploratory, we found consistent patterns across varied genotypes that persisted after stringent filtering using a conservative statistical approach (74). Furthermore, the percent of loci that exhibited a significant environmental correlation (1.8%) is well within the range of most other studies using BayEnv (74). The identified genes were functionally enriched for the Gene Ontology term, “hsa-mir-10b-5p” that corresponds to a microRNA that regulates the expression of a diverse group of genes that stimulate growth via the regulation of lipid metabolism (75). Previous studies on amphibian development have shown that lipid metabolism is dynamic; it is initially low to increase fat stores, then increases late in the larval stage to fuel metamorphosis (76). In the CTS congener *Ambystoma opacum*, increased fat stores at the time of metamorphosis significantly increases post-metamorphic survival (77), and segregating variants of the “mir-10b” gene-family may confer increased survival in short or long hydroperiod treatments, resulting in its consistent selection across larval groups. This family of miroRNAs has also been shown to promote vascular endothelial growth during the development of blood vessels (78). This function may be critical in drying ponds where oxygen content is often low (79), and increased vasculature in the gills and skin should fuel the rapid growth and developmental reorganization in both tissues that are critical for successful metamorphosis (80, 81). A recent study in the closely related *Ambystoma velasci* highlighted numerous genes that are differentially expressed during metamorphosis that are responsible for increased vascularization (82).

The protein coding gene Serum Response Factor Binding Protein 1 (“SRFB1”) was significantly correlated with hydroperiod in more than one larval group, suggesting a greater importance for this gene in mediating survival in short vs. long duration ponds. Studies on mouse model systems suggest that SRFB1 is a translational regulator that plays a significant role in cardiac aging and mitochondrial function (83). Expression of this gene has been shown to reduce mitochondrial size and oxygen consumption (84). If this functions in a similar manner in tiger salamanders (no functional studies currently exist), it may play a role in modulating the rate of metabolism in different hydroperiod regimes. Future studies that investigate the role of this gene in salamander metabolism and development through knockout (e.g. 63–65) or quantitative trait loci experiments (88) would be a valuable addition to our knowledge of its role in hybrid superiority.

Our genomic analyses provide a list of candidate genes that may underlie hydroperiod adaptation in both native and non-native hybrids. Future landscape-level studies should compare evolution in these regions to genome-wide variation in order to assess their relative importance to survival. If hydroperiod adaptation is indeed critical for survival in the California landscape, and native genotypes represent an optimal evolutionary response, then we would predict rapid fixation of native alleles at these loci. Alternatively, novel genomic combinations achieved through hybridization may have produced superior, transgressive adaptations resulting in strong selection for combinations of native and non-native alleles. Particularly if some level of hybrid superiority is deemed acceptable in a conservation framework, genome-informed management can target largely native genotypes that retain some of the fitness benefits of non-native introgression (73). Furthermore, targeting native populations that have alleles associated with short hydroperiod adaptation for conservation or translocation may help buffer CTS from future climate change and increased periods of drought (89). Further research on these candidate genes can provide the framework for making these genomically-informed management decisions.

### The Bottom Line for Conservation

Contrary to single-taxon artificial mesocosm studies (39), our naturalistic whole-pond manipulations did not detect differential selection for native or non-native genotypes associated with hydroperiods from 80 to 115 days. Assuming that this result also holds for longer hydroperiods (an assumption that also needs to be tested), reducing hydroperiod will not alter the relative success of native CTS. However, it will reduce the absolute success of any genotype, and in that sense, it remains a powerful tool if properly applied.

Reducing pond hydroperiod, particularly for key ‘source’ ponds that maintain large non-native populations within the hybrid zone, represents a relatively inexpensive strategy to reduce the overall production of hybrids without eliminating salamanders altogether from the landscape. Such an approach would have two major benefits. First, it would reduce the number and fitness of hybrid offspring, likely reducing the number of hybrid migrants that would disperse and colonize nearby ponds. And second, it allows other vernal pool functions, including the development of other endangered amphibians, crustaceans, and endemic annual plants, to continue as hybrid salamander removal strategies are developed. Recent experimental work in the CTS/BTS system has clearly shown that maintaining hybrids results in more natural vernal pool dynamics than completely removing salamanders (25). At the very least, reducing hydroperiod slows down the overproduction of hybrids and their dispersal into adjacent habitats, allowing time to enact other management strategies, including targeted hybrid removal. The critical next step for hybrid management is the development of a rapid assay to detect non-native hybrids, and key genes like SRFB1, in hours, in the field. The technology to accomplish such rapid genotyping exists (90, 91), and could be applied in combination with reduced hydroperiod to genotype, and remove, hybrids as they enter ponds to breed. Future studies should apply demographic simulations to explicitly test the extent to which hydroperiod management could affect these dynamics; our intuition is that it could be a powerful combined effort.

Extended hydroperiods also appear to have a previously unrecognized benefit to pure CTS populations. Although the ecological effects of late emergence into summer conditions on metamorph fitness are not currently known, the demographic benefits of greater size at metamorphosis are considerable and well established (40, 42). Given the effect on metamorphic size, it is likely that prolonged hydroperiods, as often characterize ponds utilized for livestock grazing, is a positive benefit for populations of pure CTS that are not at risk of non-native colonization. There are of course other ecological consequences of long hydroperiods, including their potential suitability for non-native predators ranging from mosquito fish and centrarchids to bullfrogs and crayfish, which must also be considered. However the clear benefits of prolonged hydroperiods for native CTS have never been demonstrated previously.

## Acknowledgments

We acknowledge the agencies that provided support including the USFWS, BLM, US Army, UCNRS, Burleson Consulting, and CSUMB. We thank E. Toffelmier, C. Searcy, the Shaffer Lab, and countless volunteers that made the fieldwork possible. We thank the UCLA La Kretz Center for California Conservation Science and the UCLA Stunt Ranch Reserve for partial funding.

## Notes

**Competing Interest Statement:** We have no competing interests to report.

### Competing Interest Statement

The authors have declared no competing interest.

