## Supplemental Text and Figures for "Managing invasive hybrids through habitat restoration in an endangered salamander system"

**Supplemental Information**

**Methods:**

*Pond Construction*

We designed 7 ponds in each of two sizes, large (Diameter = 9.1m, Max Depth = 69cm) and small (Diameter = 7.9m, Max Depth = 60cm), both with 15% slope. These two pond sizes were used to coarsely differentiate pond hydroperiod based on calculations using average evapotranspiration and precipitation for the site following established protocols (1). Therefore pond size is not directly considered in downstream analyses, since the hydroperiod treatment is more precise, and highly correlated with pond size. These sizes were determined using rough calculations of annual evapotranspiration rates for Monterey County. We constructed ponds following previously established designs for PVC lined ponds (1). Briefly, ponds were excavated 15cm deeper than the desired maximum depth to account for 15cm of added substrate. Basins were then lined with a 30-mil Fish-Grade aquatic safe PVC liner (FSP30 WP; Fab-Seal Industrial Liners INC.) protected above and below with layers of 16 OZ Geo Textile Padding. These three layers were installed in the basin and anchored 15cm below ground level using 12 inch (30.5cm) stainless-steel landscaping spikes with 2” (5cm) washers. Topsoil was then used to cover the liner/geotextile layers at a depth of approximately 15cm all around. These ponds were designed to fill with directly intercepted winter precipitation. We installed 48 inch (122cm) aluminum straight edge rulers attached to cinder blocks in the center of each pond to record daily water depth. We also added additional refugia from natural (branches, sticks, leaves) and artificial (cinder blocks, plywood, 15 cm PVC pipe segments) sources equally across all ponds.

In year one ponds filled November 22, 2018, larvae hatched in their source ponds approximately February 17^,^ 2019, were added to the experimental ponds around March 15 2019 and the last pond dried June 23, 2019 amounting to a maximum hydroperiod of 115 days. In year two (2020) ponds filled November 27, 2019, larvae hatched around February 4, 2020, were added to the experimental ponds March 9 2020, and the last pond dried June 3, 2020 totaling a maximum hydroperiod of 110 days. Larvae were added to the experimental ponds after they achieved sufficient size (~1.5cm) to be safely captured in a seine net and transported. Attempting to capture the larvae earlier may have imparted significant mortality and confounded the survival analyses.

*Drift Fences*

Drift fences were constructed using partially buried, 0.3 meter tall shade cloth that completely encircles each pond approximately 1m from the edge of the constructed basin. An additional line of drift fencing surrounded the entire site to ensure no hybrid salamanders escaped. Pitfall traps consisted of 1-gallon buckets buried so that they are flush with the surface of the ground and spaced every 10 meters on both sides of the drift fence. Bucket lids were modified by attaching wooden feet to the top of the lid so that the lid can be positioned over the open trap to provide shade and cover to prevent desiccation. These lids could be flipped over and used to close the traps when not in use.

*Source Pond Larvae genetic sampling*

Although sampling the true founder individuals would have been ideal, this method was not feasible for several reasons. First, the size of the tissue sample required for our target-capture protocol is too large to excise from the founder larvae without causing serious injury or death (2). Second, tissue sampling injuries would have disproportionately disadvantaged smaller larvae, potentially favoring large larvae. Third, we were unable to sequence each of the 2,730 larvae included in this study due to financial constraints, necessitating a representative sampling design. We therefore believe that sequencing a randomly drawn sample from the pool of founders, balanced across the observed size distribution, represents the most rigorous method available.

*Larval densities*

The low- and high-hybrid treatments received 120 total larvae, which is approximately 6.7 larvae per cubic meter of maximum pond volume. This density was chosen to replicate the hydroperiod mesocosm study (3) which used 6.6 larvae/m^3^. The medium-hybrid treatment received fewer total larvae across all experimental ponds (75 larvae or approximately 4.2 larvae/m^3^) due to exceptionally low breeding in wild source ponds. Given the low numbers of breeding tiger salamanders during both sampling years, we limited the total number of larvae used in the experiment to minimize our impact, particularly on the native CTS. We account for these differences by introducing larval proportion as a random effect in all applicable analyses.

*Bayesian Model Parameters*

Larval Survival

$$logit\left( p_{i} \right)= b_{HYDP}*{HYDP}_{i}+ b_{HIS}*{HIS}_{i}+ b_{0, t_{i}}$$

$$y_{i}\sim Bernoulli(p_{i})$$

The prior distribution for b_0_ was specified as a normal distribution with parameters $\mu$ and $\tau$, which were hyperpriors shared among all three treatment levels. Parameter $\mu$ was set to the logit-transformed probability of survival (p_0_) when HYDP and HIS were zero. The beta distribution was used as the prior for p_0_, with parameters $\alpha$ = 1 and $\beta$ = 1. We used a uniform distribution from 0 to 5 as the prior for $\sigma$. The value for $\tau$ was then computed as $\frac{1}{\sigma^{2}}$. A uniform prior distribution from -5 to 5 was used for both b_HYDP_ and b_HIS_. The model was iterated 500 times to explore the variation present in the larval resampling process. Each iteration used 4 independent Markov chains and 1000 iterations per chain. We combined the posterior distributions to estimate model parameters.

Metamorph Mass

$$\mu_{i}= b_{{HYDP}^{2}}*{{HYDP}_{i}}^{2}+b_{HYDP}*{HYDP}_{i}+ b_{HIS}*{HIS}_{i}+ b_{0, t_{i}}$$

$${MASS}_{i}\sim Normal(\mu_{i}, \tau)$$

The prior distribution for b_0_ was set to a normal distribution with hyperpriors mean ($\mu_{b}$) and standard deviation ($\tau_{b}$) which were shared across treatment levels. The prior for $\mu_{b}$ was set as a normal distribution with a mean of 0 and standard deviation of 0.1. $\tau_{b}$ was calculated as $\frac{1}{\sigma_{b}^{2}}$ and the prior for $\sigma_{b}$was set to a uniform distribution from 0 to 10. We used a uniform distribution from 0 to 10 as the prior for $\sigma$, $\tau$ was then computed as $\frac{1}{\sigma^{2}}$. A uniform distribution was used for all slope parameters with a range: from -30 to 0 for b_HYDP2_, from 0 to 30 for b_HYDP_, and from -10 to 10 for b_HIS_. These distributions were updated to ensure that the estimates were not restricted based on the prior.

Standardized Mass

$$\mu_{i}= b_{HYDP}*{HYDP}_{i}+ b_{GEN}*{GEN}_{i}+ b_{0, t_{i}}$$

$${log(MPL}_{i}) \sim Normal(\mu_{i}, \tau)$$

The prior distribution for b_0_ was set to a normal distribution with hyperpriors mean ($\mu_{b}$) and standard deviation ($\tau_{b}$) which were shared across treatment levels. The prior for $\mu_{b}$ was set as a normal distribution with a mean of 0 and standard deviation of 0.1. $\tau_{b}$ was calculated as $\frac{1}{\sigma_{b}^{2}}$ and the prior for $\sigma_{b}$was set to a uniform distribution from 0 to 10. We used a uniform distribution from 0 to 10 as the prior for $\sigma$, $\tau$ was then computed as $\frac{1}{\sigma^{2}}$. A uniform prior distribution from -5 to 5 was used for both b_HYDP_ and b_GEN_. A uniform distribution from 0 to 10 was applied to $\sigma$.

*Molecular Methods*

We followed a modified mybaits protocol (version 2.3.1) with our own species-specific repetitive DNA blocker c_0_t-1 (30,000 ng in 5uL in 10mM Tris-HCl, pH 8) for use in the capture reactions. Libraries were hybridized to probes for 30 hours, subjected to three high-stringency wash steps, and PCR-amplified to enriched for target DNA. Each pool was split each into 4 replicate reactions to help reduce PCR bias (4), and capture pools were then combined into two final pools (2019 and 2020 samples).

*Bioinformatics*

We first marked illumina adapters and then marked duplicate reads using picard. We used gatk to recalibrate base map-quality scores in known variant sites using a variant database from previous CTS studies (McCartney-Melstad et al., 2016; McCartney-Melstad et al. unpublished data). We used gatk HaplotypeCaller to call haplotypes over genomic regions that matched our 5,237-gene target regions (option “-L” with a Browser Extensible Data (BED) file of target regions with a 300bp buffer). These individual Genomic Variant Call Format (GVCF) files were then combined into one multi-sample GVCF using gatk CombineGVCF. We then called genotypes using gatk GenotypeGVCFs. We used gatk VariantFiltration to remove loci in the Variant Call Format (VCF) file that failed any of the following conditions: QualityByDepth (QD) < 2, MappingQuality (MQ) < 40, FisherStrand (FS) > 60, MQRankSum < -12.5, ReadPosRankSum < -8.0, QUAL < 30 (For description see URL: <https://gatk.broadinstitute.org/hc/en-us/articles/360035890471-Hard-filtering-germline-short-variants>). We used vcftools (5) to remove individual genotype calls with quality less than 20 (“--minGQ 20”) and depth less than 8 (“--minDP 8”). We also used vcftools to filter loci that failed the following conditions: were not bi-allelic (“--min-alleles 2 --max-alleles 2”), were missing data across more than 50% of individuals (“--max-missing 0.5”), had a minor allele frequency less than 10% (“--maf 0.1”). For some downstream analyses we used the “prune” plugin in bcftools (6) to filter loci that were physically linked with an r^2^ greater than 0.8 within a 1000bp sliding window (“-m 0.80 -w 1000”).

To identify diagnostic loci, we generated VCF files for the pure CTS and pure BTS separately, then filtered these files using VCFtools (“--max-maf 0.001 –max-alleles 2”) to identify loci that were monomorphic in each group. We then used custom R scripts to find loci that were fixed for different alleles in the CTS and BTS reference groups. This yielded a list of diagnostic loci with information about the species-specific origin of each allele. We used this list of loci to subset the main sample VCF such that only diagnostic loci were included. We then filtered loci using a strict 95% threshold for linkage disequilibrium using the “prune” plugin in bcftools (6) with a 1000bp sliding window (“-m 0.95 -w 1000”). This reduced the likelihood of counting two physically linked diagnostic loci, which would not represent independent data.

*Additional Metamorph Size results*

Metamorph mass was greater in 2020 than in 2019 (LMM: estimate = 4.75, CI = (3.69, 5.84), p = 2x10^-16^), which was likely a result of the lower initial larval densities used in 2020 due to limited breeding in wild source ponds.

Metamorph SVL also significantly increased with HIS (LMM: estimate = 1.66, CI = (1.38, 1.94), p = 2.0x10^-16^), although the model including both HIS and hydroperiod was not significantly different from the simpler model which only included HIS (LRT: dAIC = 0.22,$\chi^{2}$ = 2.23, p = 0.136).

**Supplemental Figures**

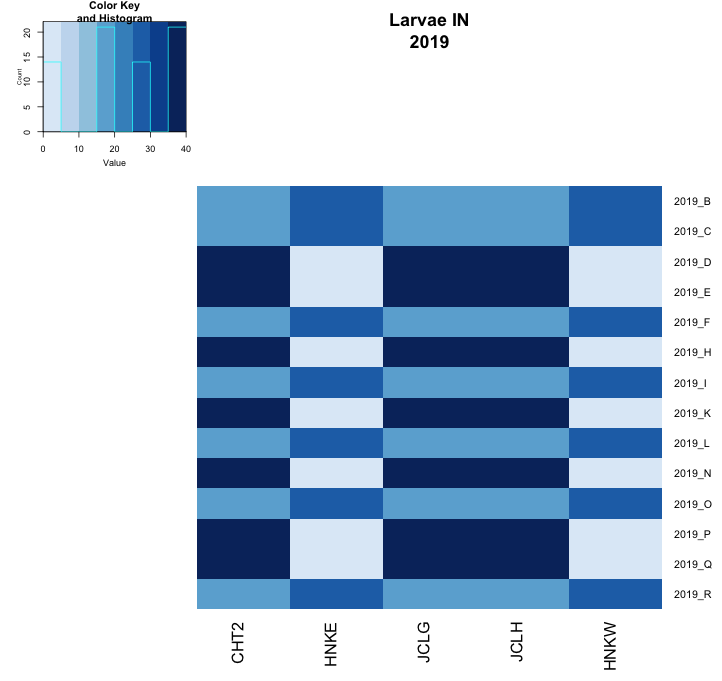

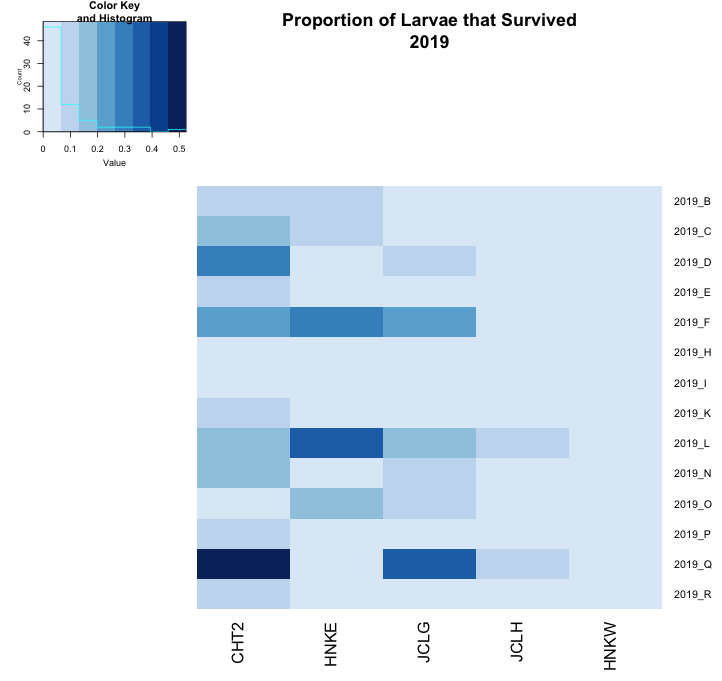

B

A

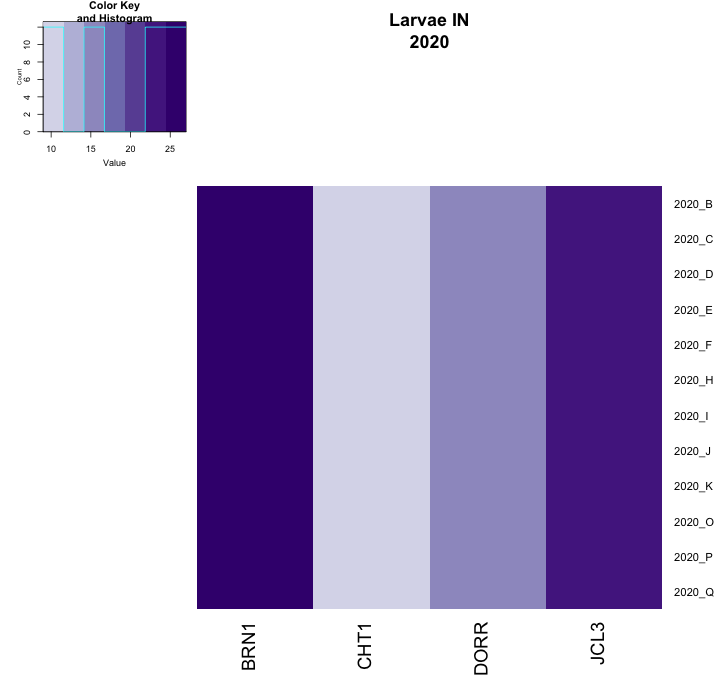

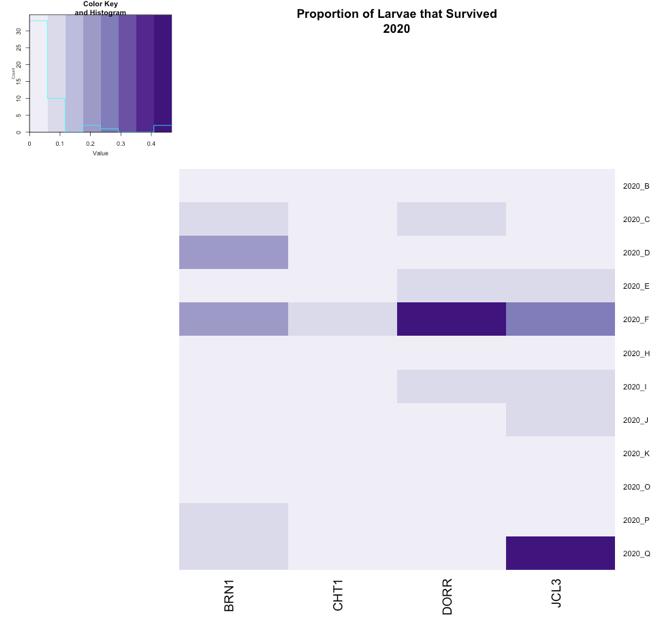

D

C

Figure SI.1: Heatmaps depicting the number of larvae from each source pond that were added to the experimental ponds (left plots, A and C) and the number of surviving metamorphs that emerged from the ponds (right plots, B and D). This demonstrates the strong dissimilarity between the starting and ending proportions. There is not a single source pond that performs exceptionally well across ponds. However, there appears to be an unequal distribution of surviving metamorphs. It appears that 1-2 source ponds make up the majority of all metamorphs that emerge from an experimental pond, suggesting a strong source pond/family group effect.

**Literature Cited (Supplemental):**

1.	T. R. Biebighauser, *Wetland restoration and construction: a technical guide* (Upper Susquehanna Coalition Horseheads, New York, USA, 2011).
